# Solute Transport in Engineered Living Materials using Bone Inspired Microscale Channel Networks

**DOI:** 10.1101/2023.07.03.547434

**Authors:** Ellen W. van Wijngaarden, Samantha Bratcher, Karl J. Lewis, Christopher J. Hernandez

**Affiliations:** Sibley School of Mechanical and Aerospace Engineering, Cornell University, Ithaca, NY 14853, USA; Meinig School of Biomedical Engineering, Cornell University, Ithaca, NY 14853, USA; Departments of Bioengineering and Therapeutic Sciences and Orthopedic Surgery, University of California, San Francisco, CA 94143; Chan Zuckerberg Biohub, San Francisco, CA 94143

**Keywords:** Engineered living materials, porous materials, bacteria, mechanical stress, microfluidics

## Abstract

Engineered living materials (ELMs) are an emerging class of materials that is synthesized and/or populated by living cells. Maintaining living cells within an ELM over prolonged periods remains a major technical challenge that limits the service life of a material. Biological materials regularly maintain robust populations of living cells. Bone maintains living cells for decades by delivering nutrients through a network of nanoscale channels punctuated by microscale pores. Nutrient transfer in bone is enabled by mechanical loading experienced during regular use. Here we identify the characteristics of channel-pore network geometries and external mechanical loading that can be used in engineered living materials to deliver nutrients to resident cell populations. Transport occurs when deformation in the microscale pore network exceeds the volume of the connecting channels. Computational models show that transport is enhanced at greater load magnitudes and lower loading frequencies and are consistent with experimental validation using microfluidic systems. Our findings provide quantitative design principles for channel-pore networks capable of delivering nutrients to materials designed to house living cells.

## 1. Introduction

Engineered living materials (ELMs) are a new class of materials that are synthesized by and/or populated by living cells. ELMs have the potential to provide novel functionalities, including self-healing, self-assembly and response to environmental stimuli ^[1–3]^. A major challenge to the development of ELMs is maintaining the viability of living cells within the material. Extending the service life of ELMs is necessary to achieve the functionalities provided by living cells beyond short-term applications.

To date, research in ELMs has focused on material synthesis by living cells or on the use of synthetic biology techniques to acheive novel functionality of the material. The most successful demonstrations of ELMs have used hydrogels which are porous enough to enable diffusion of nutrients and waste products to and from embedded cells ^[2,3,5]^. Organisms within the hydrogel can enable controlled secretion of proteins to assemble structures ^[4]^, biodegradability ^[6]^, high stretchability ^[7]^ and biocompatibility for tissue engineering ^[4,6–8]^. In all demonstrations to date, the living functionality of ELMs has been limited by the viability of resident cells, which run out of nutrients or accumulate excessive metabolic byproducts within hours of activity. Until methods of maintaining the viability of resident cells are identified, applications for ELMs will be limited to single use devices.

Naturally occurring living materials provide useful models of mechanisms of maintaining the viability of cells. Bone is a living material in which resident cells, called osteocytes, are entombed within a mineralized matrix and typically survive for decades. Osteocytes reside in pores, called lacunae (10 μm in characteristic size), connected to one another and nutrient rich surfaces through nanoscale channels, called canaliculi (0.2-0.4 μm diameter, ∼20 μm in length, Figure 1a) ^[9]^. The lacunarcanalicular channel-pore network is not subject to net fluid flow across cells, raising the question of how osteocytes receive sufficient nutrients to survive ^[10,11]^. Piekarski and Munro proposed that nutrient flow to osteocytes was enabled by dynamic mechanical stress experienced by bone tissue during regular use ^[12]^: External stresses applied to bone result in deformation of lacunae that force fluid through canaliculi into neighboring lacunae or surface nutrient sources. Upon relaxation, the lacunae expand, causing an influx of fluid containing nutrients from the neighboring source **(Figure. 1b)**. Wang and colleagues confirmed this concept computationally and demonstrated experimentally that mechanical loading to bone allows for solute transport to osteocytes that greatly exceeds what could occur with diffusion alone ^[10,12,13]^.

**Figure 1.**
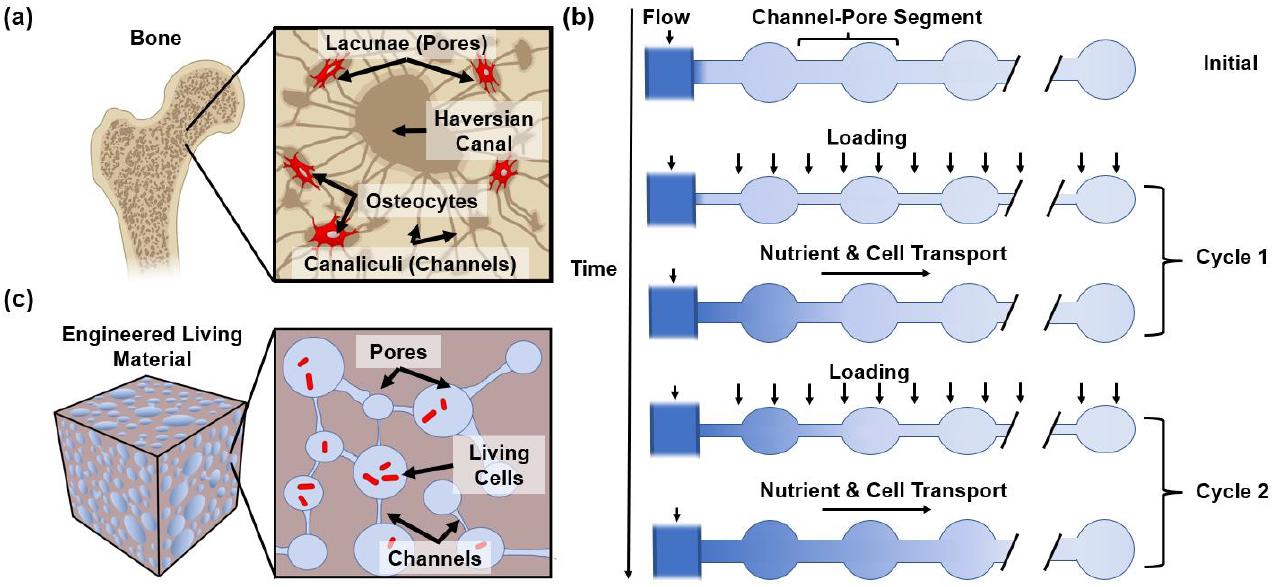
(a) Cells within bone survive in microscale pores connected to each other and nutrient rich outer surfaces through nanoscale channels. (b) A cartoon illustrating how cyclic mechanical loading can enable fluid transport within a network as described by Wang et al. ^[9]^. Each cycle of compression and expansion causes an efflux and influx of fluid from the source channel on the left, resulting in mixing and transport of nutrients within the network despite zero net fluid flow. (c) A hypothetical engineered living material in which live cells reside in pores connected to one another with nano/microscale channels.

Although the combination of channel-pore networks and cyclic mechanical loading have been shown to enable nutrient transport in bone, application of this mechanism to engineered materials requires identification of the range of possible channel-pore geometries, material substrates and forms/magnitudes of external mechanical loading sufficient to enable solute transport. An understanding of the design space for this transport mechanism would enable development of engineered living materials with longer periods of cell viability. Our long-term goal is to build engineered living materials that house embedded cells with sufficient service life for use in durable devices **(Figure 1c)**. Here we specifically ask 1) what are the channel-pore geometries that can allow mechanically enhanced solute transport to resident cells; and 2) how does the geometry and mechanical environment determine the solute distribution.

## 2. Analysis and Results

We used a computational model of two different forms of mechanically-induced solute transport to determine the characteristics of channel-pore geometries that allow nutrient transport. The models were used to determine how loading modality and geometry altered solute transport as well as the number of loading cycles required to reach a desired solute concentration in each pore and the relationship between the distance from the nutrient source and time to reach a desired solute concentration. The computational results were validated experimentally using microfluidic systems.

### 2.1. Theory

Here we examine a network consisting of thin channels punctuated by larger pores as is seen in osteocyte networks in bone (Figure 1a). Fluid may only enter or exit the channel-pore system through the connection to the source, i.e., the inlet is the only outlet and there is no net convective flow through the system. Such a geometry is easily applied to ELMs in which the channel-pore network is created by a surface treatment or is due to stochastic processes during manufacturing (i.e. an explicit network is not required). Although the entire channel-pore network may be complex (Fig 1a, c), Wang et al. have shown that transport mechanisms in the complex network seen in bone is adequately modeled by a linear connection of channel-pore segments ^[9]^ (Figure 1b). We likewise assume that the fluid in the channel-pore system is Newtonian and that the length scales are small enough to assume laminar flow, i.e. length scales approximately as small as 10 nm and below 1mm with low Reynolds numbers ^[14]^.

Although diffusion may occur within the channel-pore network, more rapid solute transport is generated when the channel-pore system is deformed by cyclic mechanical loading. Wang et al. illustrated the concept in bone using cyclic compressive mechanical stress which generates compressive volumetric strain within the pores, forcing fluid out of the pore through the channels. When the applied mechanical stress is removed, the pore regains its initial volume and there is an influx of fluid from the source. No net fluid exchange occurs over the course of a full cycle since the volume of fluid leaving the pore is the same as that returning upon unloading. However, if the fluid displacement from the pore exceeds the channel length, L, solute transport can occur upon unloading because some of the fluid that returns to the pore has mixed with the source (or preceding pore). Wang et al. found that the fluid volume leaving a pore, *q*_*i*_, is equal to the change in volume of the pore *ΔV*. The fluid volume leaving the pore is also related to the displacement of the fluid down the channel, *d*_*i*_ and the cross-sectional area of the channel, A_ch_(or total cross-sectional area of all connecting channels) as shown below ^[9]^:

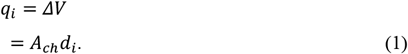

For mixing and solute transport to occur, the displacement must exceed the channel length. Similarly, the volume of fluid moved from the source (or preceding pore) to the pore upon unloading is the difference between the volume of fluid displaced and the volume of the channel, *V*_*c*_:

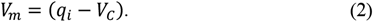

Solute concentration in a pore located *i* segments from the source is given by Equation 3. Assuming that the influx, *q*_*i*_, and efflux, *q*_*i*+1_, between pores is equal simplifies to Equation 4 (see Supporting Materials S2.1 for a complete derivation):

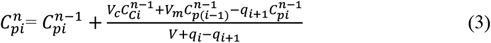

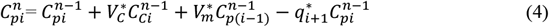

where 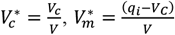 and 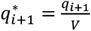 are non-dimensional parameters. 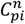 represents the solute concentration at pore *i* after *n* number of cycles. Similarly, 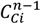 represents the concentration in the channel at *n* − 1 cycles, *V*_*c*_ represents the average channel volume, *V* is the pore volume and q_*i*_ is the flow leaving pore *i* − 1 and moving toward pore *i*. To solve for the solute concentrations in each pore for each cycle of loading, Wang and colleagues solved Equation 3 iteratively. A similar approach is taken in the current work.

Notably, in a situation where pore volume and channel volume are approximately uniform in size, the relationship provided by Equation 3 can be simplified to express the change in solute concentration caused by one cycle of loading as follows:

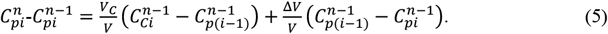

Equation 5 presents two terms, the first associated with channel-pore geometry 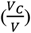, and the second, determined by the magnitude of volumetric strain caused by mechanical loading 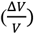. If the amount of solute required to maintain cell viability within a time period is known, Equation 5 can be used to estimate the maximum distance from a source (in channel-pore segments) that can house viable cells.

### 2.2. Mechanically Enhanced Solute Transport Caused by Cyclic Mechanical Stress

Here we consider deformations of the channel-pore network caused by mechanical stresses applied to the bulk material **(Figure 2a, b)**. The volumetric strain (*ΔV/V*) in the pore caused by the applied mechanical stress is related to the bulk modulus by:

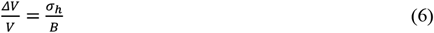

where *σ*_*h*_ is the hydrostatic stress and *B* is the material bulk modulus, defined in an isotropic material as:

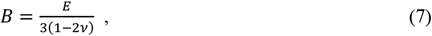

in which *E* is Young’s modulus and *v* is Poisson’s ratio.

**Figure 2.**
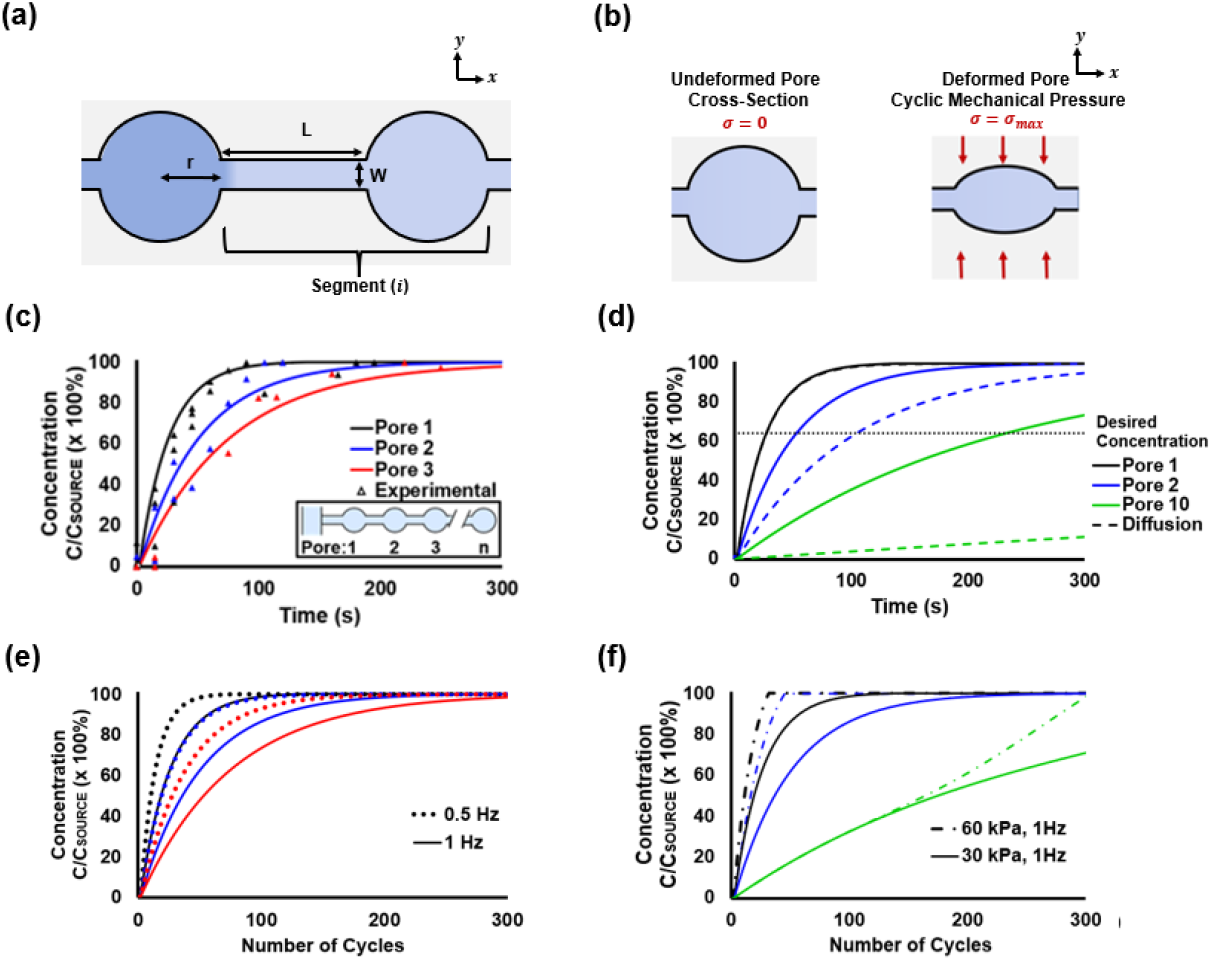
(a) A single segment of the channel pore network is shown. (b) Cyclic compression deforms pores causing mixing of the fluid between pores. (c) The concentration of the dye within the first three pores of the channel-pore network during cyclic loading is shown. The lines indicating the computational models and the data points showing the experimental results (loading condition shown: 30 kPa and 1 Hz). Colors indicate pore number away from the source. (d) Computational models show the solute concentration in pores is greater when cyclic mechanical loads are applied to the material (solid line) than would be expected from diffusion alone (dashed lines). (e) Decreasing the frequency of the applied loading enhances solute transport (loading condition shown: 30 kPa). (f) Increasing the loading stress by an order of magnitude only slightly enhances solute transport, consistent with the findings of Wang et al. in bone.

Substituting Equations 1 and 5 with the condition that fluid displacement, *d*_*i*_ must exceed channel length L, leads to the condition under which cyclic mechanical loading enhances solute transport:

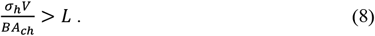

A computational model was created to apply Equation 3 iteratively for each pore at each loading cycle to find the solute transport between pores. The model was used to predict pore solute concentration at different distances from the source, compare the results to diffusion and evaluate the effects of cyclic loading frequency and magnitude.

Cyclic mechanical loading increased solute concentration over time, initially at pores closest to the source but eventually at pores further away **(Figure 2c)**. To confirm the accuracy of the computational model, a microfluidic system with a pore network was manufactured from polydimethylsiloxane (see Methods). The microfluidic system was wetted with deionized water, then a fluorescent solute was flowed through the source channel (see Methods) and the fluorescence was measured in pores. Changes in solute concentration within the pores were measured by fluorescence intensity. The experimental findings closely matched the results of the computational model both during cyclic mechanical loading and for the diffusion limited amounts of solute transport that occur without cyclic loading (**Figure 2c**).

The computational models demonstrate that cyclic loading greatly enhances solute transport within the system. Constant flow of the solute solution through the source or insufficient deformation of the pores during loading (see Equation 8) only allows for transport via diffusion of the fluorescent tracer into the channel-pore network **(Figure 2d)**. However, cyclic mechanical loading that causes sufficient deformations of the pores results in transport much greater than expected from diffusion alone (**Figure 2d**). Diffusion delivers solute to the pore closest to the source at a rate similar to that acheived with cyclic mechanical loading. However, only small amounts of solute reached the tenth pore in the system over the time period simulated. The minimum solute delivery required to maintain cell viability will depend on the solute, type of cell (species and strain), cell metabolism and toxicity of cell metabolic byproducts. As an example, we consider a desired solute concentration of 65% of the source (horizontal dotted line in Figure 2d) (see justification in Supplementary Material 2.2). The transport of solute per cycle can be enhanced by decreasing the loading frequency **(Figure 2e)**, although small increases can also be achieved by increasing the loading magnitude **(Figure 2f)**.

### 2.3. Mechanically Enhanced Solute Transport Caused by Cyclic Fluid Pressure

Here we consider the situation where the walls of the channel-pore network have low enough stiffness such that cyclic fluid pressure is sufficient to generate volumetric strains within the pores to enhance solute transport **(Figure 3a)**. In this situation, fluid pressure within the channel-pore system varies with distance from the source and therefore a computational model integrating hydraulic circuit calculations is required (Supporting Information sections 1.1.-1.3.). The hydraulic resistance of each channel and pore was determined using hydraulic circuit calculations considering the channel length, width and pore radius (Figure 2a, Supporting Information 1.1.)^[15]^. The resistance of the inlet channel and the applied flow rate were used to determine the pressure at the entrance to the channel-pore system. The estimated change in pressure across each segment determined with hydraulic circuit calculations was verified using a COMSOL flow simulation **(Figure S1)**.

**Figure 3.**
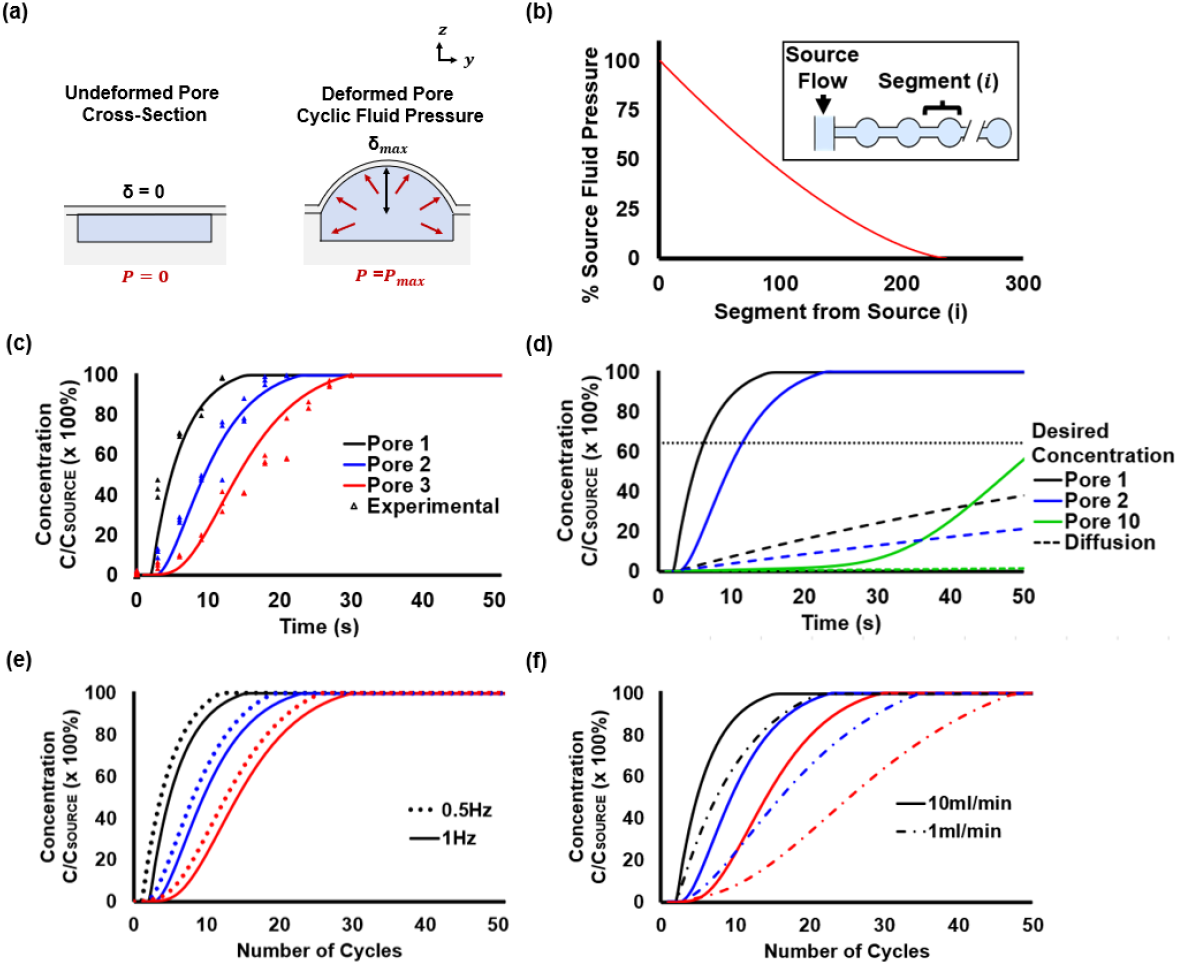
(a) Cyclic fluid flow that causes deformation of channel walls can also enhance transport. (b) Pressure declines due to hydraulic resistance. (c) Experimental results closely match the theoretical model. Cyclic fluid flow in a deformable channel-pore system allows for complete mixing (C/C_0_=100%) (peak flow rate of 10 ml/min at 1 Hz). The time for the pore to reach saturation is substantially greater when for pores farther from the source. (d) The resultant concentration enhanced by cyclic fluid flow (solid lines) rapidly exceeds the minimum nutrient distribution, but diffusion alone is insufficient (dashed line). (e) Decreasing the loading frequency increases solute transport per cycle. (f) Increases the loading magnitude enhances solute transport per cycle with the effect more noticeable for pores farther from the source.

A cyclic fluid flow at the inlet generates cyclic pressure within the source channel resulting in pressure increases within the channel-pore network. If the walls of the channel-pore system are sufficiently compliant, cyclic pressure causes expansion of the channel-pore system to allow fluid to flow from the source channel into the channel-pore system. Unlike the mechanical stress state, however, fluid pressure declines as fluid passes through channels and pores due to hydraulic resistance. The reduction in pressure across each channel segment results in lower pore pressure further away from the source and therefore smaller deformation in pores further from the source channel. As a result, pores far from the source do not deform sufficiently to allow for fluid mixing **(Figure 3b)**.

The magnitude of cyclic fluid pressure enhances solute transport. The models closely matched experiments using microfluidic pore-networks (Figure 3b). Larger magnitude loading result in a greater volume of solute influx for each cycle, increasing the concentration of solute in the pores more quickly **(Figure 3c)**. The applied cyclic load enhances transport, well beyond what is achieved through diffusion alone **(Figure 3d)**. The effect of cyclic loading on solute concentration is more pronounced at pores farther from the source channel (if they deform sufficiently to allow mixing, see pores 2 and 10 in Figure 3d). Decreasing the frequency reduces the number of cycles for the fluid to reach saturation concentration as there is additional time for diffusion to act **(Figure 3e)**. Similarly, the loading magnitude can be increased to achieve greater volumetric strain in the channel, increasing the pore concentration more quickly **(Figure 3f)**.

### 2.4. Relation between Depth of Solute Transport to Geometry and Loading

Our analysis makes it possible to estimate the distance from a nutrient source at which resident cells within the system will receive a specified concentration of nutrients over a period of time. Equation 5 highlights *V*_*c*_*/V* as a key parameter in determining the solute concentration of a pore at any distance from the source channel and is a useful parameter for design. The maximum distance that acheives a desired solute concentration over a specified time period can be found by determining which pores have attained a desired solute concentration. Here we present the relationship between channel-pore geometry and maximum distance at which a desired solute concentration is achieved. Specifically we consider a time period of one hour at 30 kPa or 1 ml/min respectively **(Figure 4a and b)**. In both cyclic mechanical loading and cyclic fluid pressure the relationship between geometry and solute concentration can be approximated with a power law (inset).

**Figure 4.**
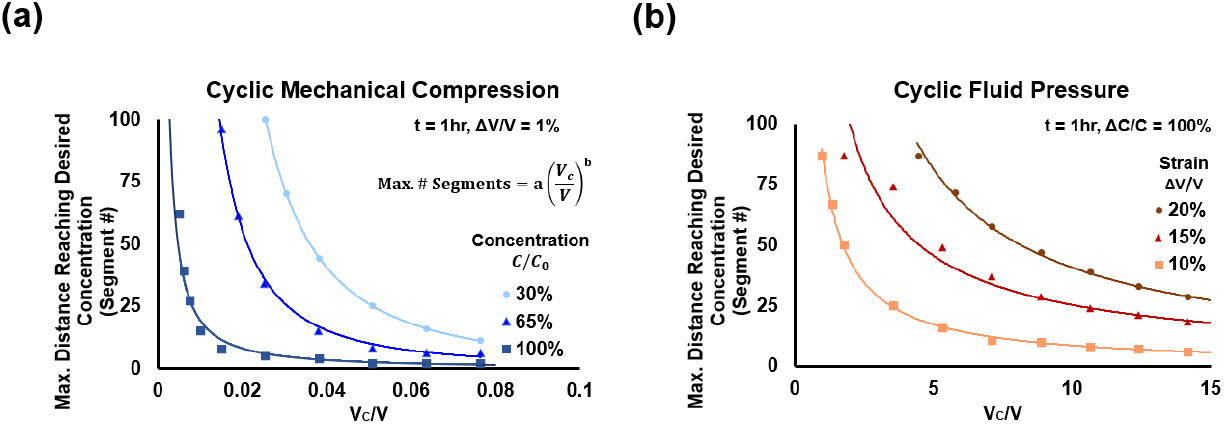
The relationship between channel pore geometry (*V*_*c*_*/V*) and the number of channel pore segments away from the source required to reach a desired pore concentration is shown. Constants a and b differ based on the volumetric strain and the desired concentration for (a) cyclic mechanical compression (volumetric strain = 1%) and (b) cyclic fluid pressure showing the maximum pore receiving solute. These curves can be applied to determine the maximum distance at which a pore concentration is estimated to be able to support a viable cell population.

The ratio of *V*_*c*_*/V* is key to the solute transport for both loading modalities, however various differences exist. As mechanical compression applies deformations throughout the pore network the change in concentration is gradual and the lines for different concentrations can be displayed for a given time point, as shown in Figure 4a. In contrast, cyclic fluid flow can introduce sharp changes in concentration as the pressure losses between pores result in different volumes of fluid being delievered to the pores for each cycle. Instead, the volumetric strain can be adjusted in conjunction with the geometry to acieve difference desired maximum distances values for solute delivery (Figure 4b).

## 3. Discussion

Our analysis estimates solute transport within a channel-pore system using design traits (channel-pore geometry), habitual mechanical use (cyclic mechanical or fluid pressure) and the desired time or number of cycles for solute transport to resident cells. We find that solute transport can occur when the change in volume of the pores exceeds that of the connecting channels. The changes in pore concentration are dominated by the ratio of channel volume to pore volume (*V*_*c*_*/V*) and the volumetric strain experienced by the pores (*ΔV/V*) during mechanical stimulation. Therefore, altering the geometrical ratio of the channel and pore volume and ncreasing the loading magnitude to apply a higher volumetric strain introduces additional solute into the pores. Lastly, decreasing the loading freqency provides additional time for diffusion over each cycle resulting in a small enhancement of trasnport. These findings provide a baseline criteria for the design of channel-pore systems to house resident cells inside impermeable matrices.

Our solution to the challenge of solute delivery within a solid, impermeable matrix requires cyclic loading (either mechanical or through fluid pressure). Mechanical loading is commonly applied to engineering materials during use and is therefore present in many situations in which engineered living materials could be applied. While this requirement can enable greater solute transport and potentially longer periods of cell viability within the ELM, a disadvantage of this requirement is that solute transport would be severely reduced if the cyclic mechanical loading was suspended (for example a long period of disuse). Despite this limitation, the channel-pore system we have described has the potential to increase the service life of resident cells well beyond that possible with current systems that are limited by diffusion. As mentioned above, we have used a simple model system in which channel-pore systems are created with a high degree of precision (the experimental model created with microfluidic designs). In practice, channel-pore systems would more likely be fabricated in a stochastic fashion, characterized with an average pore volume and average channel dimensions. Additionally, our findings illustrate some limitations regarding the distance a solute can be distributed away from the source (Figure 4a and b). Engineered living materials using this solute delivery approach will therefore be limited by the surface area of the solid that is in contact with source fluid. In situations where a greater number of resident cells is desired, a structure with increased surface area (for example the use of a lattice) might increase the population of living cells.

Although our analysis presents an attractive new design strategy, there are some limitations in our analysis that must be considered. First, our approach focuses on channel dimensions that satisfy assumptions used in micro/ nanofluidic systems with dimensions in a micro/nanofluidic range, approximately between 10 nm and 1mm. This assumption is reasonable for channel-pore systems that are well approximated with laminar flow, but are not so useful for larger scale channels in which inertial forces are considerable. Secondly, our analysis has assumed that stress relaxation and creep are negligible during the loading frequencies considered. Viscoelastic or poroelastic analyses would be required for materials/loading in which these properties are relevant. Lastly, in practice the channel-pore systems will also have living cells and extracellular material including the constituents of biofilms or extracellular matrix. While the presence of cells and other materials within the channel-pore system may reduce transport by reducing fluid volumes in the channel-pore system, the basic concepts shown here will still be valid (as confirmed by Wang and colleagues ^[9]^). Despite these limitations, the results of the computational models closely match experimental results, suggesting that the computational model encompasses the major contributors to solute transport.

## 4. Conclusions

Living cells within engineered living materials have so far been limited to permeable substrates such as hydrogels and/or only a few hours of living cell activity. Our findings suggest that viability may be enhanced in applications of cyclic loads. Our analysis demonstrates how nutrients might be delivered to living cells embedded within the ELM, allowing the living component of an ELM to be located farther from a nutrient source, thereby increasing the population of living cells within the material and also allowing for living cell functions (sensing, repair) further from external surfaces. Including design aspects in ELMs, that enhance viability of the cells, has the potential to enable more rapid development and application of this new class of materials.

## 5. Experimental Methods

### 5.1. Microfluidic Device Design

#### Microfluidic Device Design

We created microfluidic systems to test channel/pore geometries, and magnitude and frequency of mechanical loading/flow. Each device consisted of a source channel with inlet and outlet, a linear channel/pore network and an exit from the channel/pore network to facilitate wetting. A total of 10 devices were fabricated with ranges in channel length (50 μm to 200 μm), pore radius (300 μm to 2500 μm) (Table S1).

#### Microfluidic Device Fabrication

Hydraulic circuit calculations for the pore network geometries were performed using MATLAB (2022b, MathWorks, USA). The channel-pore networks were drafted using SolidWorks software (SolidWorks Corp., USA) and a negative mold was manufactured from VeroBlue using a high resolution stereolithography on an Objet 30 Scholar printer (Stratasys, Israel) using a high-resolution finish and mold release spray (Mann, USA) to enable polymer casting.

Printed molds were thoroughly cleaned with soap and deionized water. Device dimensions were confirmed using an inverted spinning disk confocal microscope (Olympus IX-83, Tokyo, Japan). A 184 SYLGARD elastomeric kit (Ellsworth Adhesives, USA) mixed at a 10:1 ratio of polydimethylsiloxane (PDMS) base to a curing agent was degassed in a vacuum chamber for 30 minutes and poured into the mold. The device was allowed to cure for 6 hours at 50 °C ^[16]^. The PDMS layer was then removed from the mold and plasma bonded to a 200 μm PDMS membrane to enable imaging of the pores. The membrane layer was made by spin coating and curing PDMS on a transparency sheet (WS-400A spin coater, Laurell Technologies, USA). Bonding was completed using a plasma bonder with room air (PDC-001-HP, Harrick Plasma, USA). Tubing, with a 1/16 in inner diameter, and luer lock connectors interfaced the device to a syringe pump (McMaster Carr, USA). The final device channel width, length and pore diameter were confirmed by observation with a microscope.

#### Fluid Displacement due to Cyclic Mechanical Loading

The microfluidic device was secured in custom fixtures beneath an upright microscope **(Figure 5a-d)**. Fixture alignment and microfluidic device dimensions ensured that the PDMS device would not substantially bend (see Supplementary Information section 4.0).

**Figure 5.**
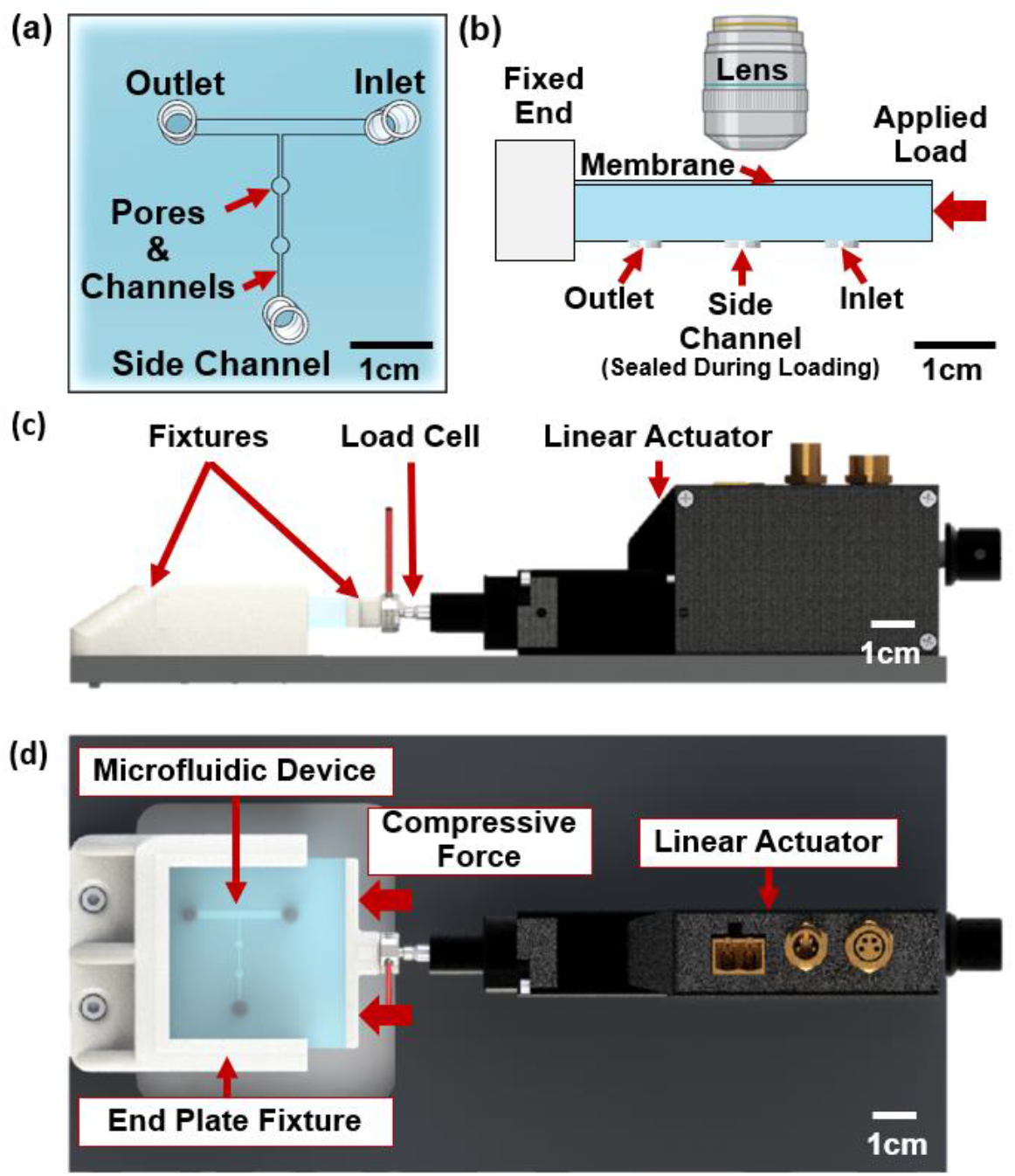
(a) The microfluidic device design consisting of an inlet, outlet and side channel with channel-pore segments. The number of side channels, pores, pore radii, and inter pore length differed between designs tested (Table S1). (b) Side view diagram of the microfluidic device is shown. The left side is fixed while a distributed load is applied to the right side. The inlet and outlet of the device are opposite the microscope objective. Imaging during mechanical loading was conducted using an upright microscope for mechanical testing. (c) The side view of the custom mechanical testing setup showing the plate transferring force from the linear actuator, positioned on the right, via the 20 N load cell and linear actuator. (d) Top view is shown depicting the placement of the microfluidic device, end plate fixture and baseplate supporting the components.

Mechanical loading was applied to the devices using a custom-built microtester (Figure 5 c, d), that included a micro linear actuator (X-NA08A25-S), Zaber Technologies, Vancouver, Canada). A haversine waveform was applied to the PDMS device using the actuator. Fluorescent images of pores within the microfluidic device were collected at the completion of a loading cycle to ensure the same field of view for each image. Custom supports were created from polylactic acid (PLA) to support the microfluidic device and enable one-dimensional loading. The supports and linear actuator were mounted on a ¼ in thick aluminum plate, that included a hole for tubing and imaging. A LCM100 micro in-line load cell (FSH04398, FUTEK Advanced Sensor Technology, Inc., Irvine, CA, USA) measured the applied force. The system was controlled in a closed loop via a custom LabVIEW program (National Instruments Corp., TX, USA). After wetting the channel-pore network the exit tot he channel-pore system was sealed so that the source channel was the only way for fluid to enter or exit the channels.

#### Fluid Displacement due to Cyclic Fluid Flow

Cyclic fluid flow was applied to the device using a computer-controlled syringe pump (Fusion 400, Chemyx, Stafford, TX). The application of cyclic fluid flow resulted in cyclic pressure within the source channel. The PDMS layer covering the channels was relatively thin (250.5 ± 71.5 μm) so that fluctuations in pressure would deform the layer, changing the volume of the pore. The resulting increases in fluid pressure are sufficient to noticeably deform the walls of the PDMS device, thereby generating the volumetric strain required for transport of solute from one pore to another. However, unlike mechanical loading, pore deformations downstream from the source are smaller as a result of pressure losses due to the hydraulic resistance in the channels.

The volumetric strain in each channel segment was determined from the deflection found using membrane plate theory (Supporting Information 1.1.). The primary volumetric change in the pores occurred from deflection of the thin layer of PDMS on the top of the device (Fig. 3a). Although the channels are also deforming during application of cyclic fluid pressure, the small size of the channels (and corresponding small deformations of the overlying layer of PDMS) lead to only negligible changes in channel cross-section and hydraulic resistance.

The hydraulic resistance for each channel-pore segment was determined using a hydraulic circuit model to find the pressure difference between each pore. The distance fluid traveled over each half cycle was then determined (Supporting Information 1.3. Table S1 for details regarding each individual design considered).

#### Imaging Protocol

Cyclic fluid flow and diffusion tests were conducted using an inverted spinning disk confocal microscope (Olympus IX-83, Tokyo, Japan). A 40× air objective lens with a 488 nm laser was used for fluorescent imaging. 1 mM Calcein was used as a solute to measure the changes in concentration in pores.

An upright, two photon microscope (Bergamo II, Thorlabs, Newton, NJ) was used for mechanical tests with a 20× water objective lens. All images were collected using a 512 × 512-pixel array using 50% of the maximum laser power, 100mW, operating at an excitation wavelength of 920 nm (Chameleon Discovery NX TPC, Coherent, CA, USA).

## Supporting Information

Supporting Information is available from the Wiley Online Library or from the author.

## Acknowledgements

NSF 2055214, 2135586 (C.J.H.), This work was performed in part at the Cornell NanoScale Facility, a member of the National Nanotechnology Coordinated 68 Infrastructure (NNCI), which is supported by the National Science Foundation (Grant NNCI69 2025233).

Imaging data was acquired through the Cornell Institute of Biotechnology’s Imaging Facility, with NIH 1S10OD010605 funding for the shared Andor Revolution Spinning Disk Confocal Microscope.

## Solute Transport in Porous Materials using Bone Inspired Microscale Channel Networks

*Figure:*

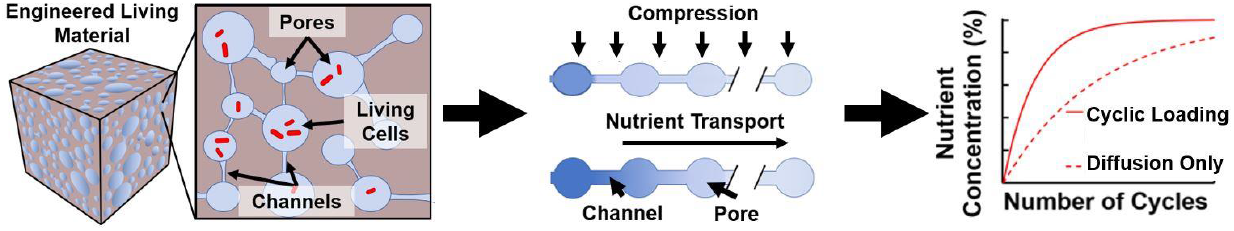

*Short Text:* Micro/nanoscale channels punctuated by pores can allow distribution of solutes to cells deep within a material without net flow through. Solute transport is enhanced by cyclic mechanical loading during regular use, far exceeding that caused by diffusion.

