## Supporting Information for "Solute Transport in Engineered Living Materials using Bone Inspired Microscale Channel Networks"

### S1.0. Model Overview

### S1.1. Cyclic Fluid Pressure in a Deformable Channel-Pore System

The application of cyclic fluid flow to a source channel will result in cyclic fluid pressure at the entrance to the channel-pore network. The channel cross-section was modeled as rectangular, resulting in a hydraulic resistance of:

$$R_{h_{0\,channel}} = \frac{12\mu L}{w h_0^3 (1 - \frac{0.63 h_0}{w})} \tag{S1}$$

where $\mu$ is the dynamic viscosity of the fluid (approximated as water with a value of $8.9\ x\ 10^{-4}$ Pa s), $L$ is the channel length, $w$ is the channel width, and $h_0$ is the channel height.

The thin membrane sealing the channel and pore deflects when a cyclic pressure is applied (Figure 3a in the main document). When pressure is applied the volume of the channel is increased and the hydraulic resistance is decreased. The average channel resistance was determined by estimating the deformed channel height as the original height plus half of the deflection.

The cyclic fluid pressure, $p$, was applied as a sinusoidal waveform:

$$p = \frac{-p_1}{2} \cos \omega t + \frac{p_1}{2} \tag{S2}$$

where $\omega$ is the angular frequency, $t$ is time and $p_1$ is the peak fluid pressure at the channel entrance,

$$p_1 = \frac{12\mu L Q}{w h^3}. \tag{S3}$$

where $Q$ is the flowrate. The membrane deflection at the center of the rectangular channel, constrained on both sides, is found from plate theory to be[1]:

$$\delta = \frac{w}{8} \left[ \frac{24(1-\nu^2) p w}{E d} \right]^{\frac{1}{3}}, \tag{S4}$$

where $E$ is the Young's modulus which for PDMS ($1.32\ x\ 10^6$ Pa [2]), $d = 2.5\ x\ 10^{-4}$ m is the membrane thickness, and $\nu = 0.499$ is Poisson's ratio [2]. End effects could be neglected as the length to width ratio exceeds 5 [3]. The new channel height could be approximated using the original height plus half of the deflection

$$h = h_0 + \frac{\delta}{2}. \tag{S5}$$

The deformed channel volume was approximated using the deformed height and assuming a constant rectangular cross-section. The deformed channel resistance, $R_{h_{channel}}$, was found by replacing $h_0$ in Equation S1 by Equation S5.

Hydraulic circuit calculations determined the fluid pressure at the entrance to the channel pore system (Equation S2). The pressure in subsequent channel segments was found iteratively using the hydraulic resistance (Equation S1) and the pressure at the entrance of the side channel.

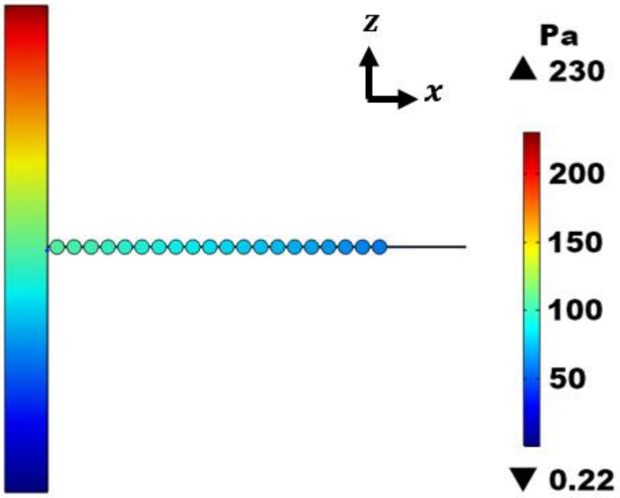


**Figure S1:** The Pressure contour map of a fluid influx indicates a drop in the pressure along the side channels with the pores primarily at constant pressure as indicated by a uniform color, (Inlet flowrate of 10 ml/min).

The hydraulic resistance of the undeformed pore is given by

$$R_{h_{0_{pore}}} = 2\int_{x=0}^{x=r} \frac{12\mu dx}{wh^3\left(1-\frac{0.63h_{pore}}{w}\right)} \qquad \text{(S6)}$$

where $r$ is the pore radius, $x$ is the cross-sectional position along the pore and $w$ is equal to the width at the cross-section position $x$. The width, $w$, is twice the value of R, the distance to the edge of the pore at position x, as illustrated in **Figure S2** or:

$$w = 2\sqrt{r^2 - x^2}. \qquad \text{(S7)}$$

The deformation of the thin layer at the top of the pore during cyclic pressure was determined from the pore pressure using plate theory[1] The resulting deflection of the pore surface modifies the hydraulic resistance of a pore in the channel network shown in Figure S2b. The maximum deflection at the center is $\delta_{max}$ is defined as:

$$\delta_{max} = 0.662R\ \sqrt[3]{\frac{pR}{Ed}}\ . \tag{S8}$$

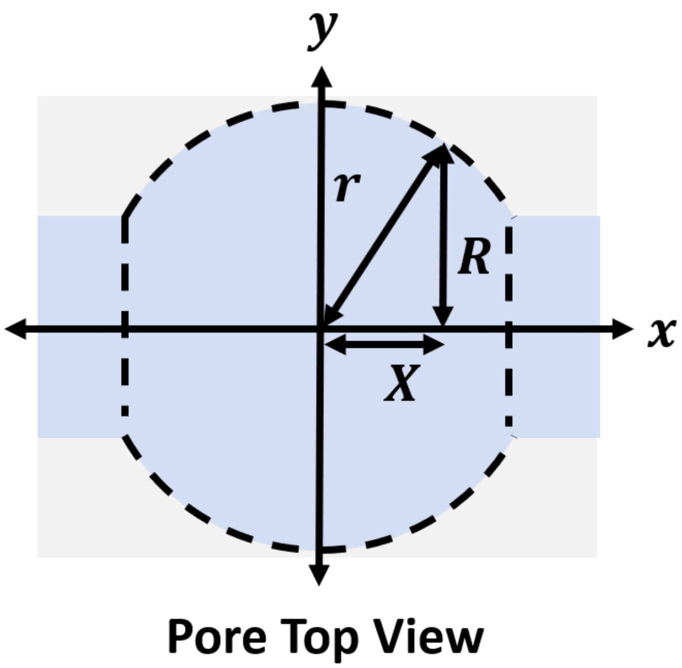


**Figure S2.** Top view of a pore is shown illustrating variables used in Equations S6-S8.

The deformed surface of the PDMS forms a paraboloid. Since the volume of a paraboloid is half that of a cylinder of the same height, the maximum deflection could be divided by two to determine the average paraboloid height, $h_{pore}$, for the deformed pore. This deformed height was substituted into the integral in Equation S6 to estimate the new deformed pore resistance and volume. The total deflected channel and pore resistances and volumes were added to determine the change in resistance and volume over a half pressure cycle to determine solute transport. Pore network geometries for designs simulated in this study are summarized in **Table S1**.

**Table S1.** Geometry of tested pore network designs is shown. W, L and R refer to the width, length and radius of the designs, indicating which dimension the design was analyzing. Note that Design Cyclic Flow: W2 was also used as Cyclic Flow: L2 and Cyclic Flow: R1 to compare the changes in channel length and pore radius to reduce the number of devices required. This provided three data points for comparing the changes in width, length and radius respectively.

| Design | Channel Width (µm) | Channel Length (µm) | Pore Radius (µm) | Min. Cyclic Compressive Stress for Solute Transport (Pa) |
|---|---|---|---|---|
| Cyclic Flow: Baseline | 50 | 100 | 300 | 48655 |
| Cyclic Flow: Width 2 | 100 | 100 | 300 | 194621 |
| Cyclic Flow: Width 3 | 150 | 100 | 300 | 437898 |
| Cyclic Flow: Length 1 | 100 | 50 | 300 | 97310 |
| Cyclic Flow: Length 3 | 100 | 200 | 300 | 389242 |
| Cyclic Flow: Radius 2 | 100 | 100 | 500 | 42038 |
| Cyclic Flow: Radius 3 | 100 | 100 | 750 | 12455 |
| Cyclic Mech.: 1 | 50 | 50 | 2000 | 82 |
| Cyclic Mech.: 2 | 50 | 50 | 2500 | 42 |
| Cyclic Mech.: 3 | 75 | 50 | 2500 | 95 |

**S1.2. Total Hydraulic Resistance**

The hydraulic resistance of a segment of the system at a given time, $R_{hyd}(t)$, was found by summing the values for the channel, $R_{channel}(t)$, and the pore, $R_{pore}(t)$.

$$R_{hyd}(t) = R_{pore}(t) + R_{channel}(t). \tag{S9}$$

The channel-pore resistance decreased by an average of 13.2% in the cyclic flow designs (Table S1) during each fluid cycle. The values for the hydraulic resistance and the volume were determined for both zero deflection and max deflection **(Table S2).**

The average hydraulic resistance, $R_{hyd\ ave}$, was used to estimate the reduction in pressure along the channel due to cyclic fluid flow using the values presented in Table S1. The hydraulic resistance is the average of the hydraulic resistances for the undeformed, $R_{hyd\ undef}$, and deformed, $R_{hyd\ def}$, cases,

$$R_{hyd} = \frac{(R_{hyd\ undef} + R_{hyd\ def})}{2} \quad . \tag{S10}$$

The change in fluid pressure, $\Delta p$ across the channel-pore segment, was calculated using the average hydraulic resistance and the average flowrate, $Q_{ave}$, as:

$$\Delta p = -R_{hyd\ ave} Q_{ave} \quad . \tag{S11}$$

**Table S2.** Volumes and hydraulic resistances a single channel-pore segment for a cyclic flow rate of 10 ml/min, 1 Hz. The results for the mechanical design geometries under cyclic flow loading are also provided to demonstrate how the hydraulic resistances can vary due to dimension changes.

| **Design** | **Hydraulic Resistance (Pa s/m$^3$)** | **Hydraulic Resistance at Peak Loading (Pa s/m$^3$)** | **Volume (μm$^3$)** | **Volume at Peak Loading (μm$^3$)** |
|---|---|---|---|---|
| Cyclic Flow: W1 | 6.08E+11 | 5.42E+11 | 1.44E+7 | 1.73E+7 |
| Cyclic Flow: W2 | 4.78E+10 | 4.27E+10 | 2.92E+7 | 3.21E+7 |
| Cyclic Flow: W3 | 1.14E+10 | 1.03E+10 | 4.44E+7 | 4.73E+7 |
| Cyclic Flow: L2 | 3.33E+10 | 2.84E+10 | 2.87E+7 | 3.16E+7 |
| Cyclic Flow: L3 | 7.66E+10 | 7.14E+10 | 3.02E+7 | 3.31E+7 |
| Cyclic Flow: R2 | 4.73E+10 | 3.92E+10 | 7.95E+7 | 9.57E+7 |
| Cyclic Flow: R3 | 4.69E+10 | 3.60E+10 | 1.78E+8 | 2.40E+8 |
| Cyclic Mech.: 1 | 3.68E+11 | 2.38E+11 | 9.40E+9 | 2.49E+10 |
| Cyclic Mech.: 2 | 3.68E+11 | 2.38E+11 | 1.47E+10 | 3.88E+10 |
| Cyclic Mech.: 3 | 8.64E+10 | 5.00E+10 | 1.47E+10 | 3.08E+10 |

#### S1.3. Fluid Displacement caused by Channel and Pore Deflection

The volumetric strain, $\Delta V/V$ was found by dividing the deformed total volume of the pore and the channel by the original volume:

$$\frac{\Delta V}{V_0} = \frac{(whL+\pi r^2 h_{pore})-(wh_0L+\pi r^2 h_0)}{(wh_0L+\pi r^2 h_0)}. \tag{S12}$$

The average flow rate over half a cycle, $t_{half}$, is then:

$$Q_{ave} = \frac{\Delta V}{t_{half}} \tag{S13}$$

$$= -\frac{\frac{\Delta V}{2}\cos(\omega t)+V_0}{t_{half}}.$$

The fluid velocity, $u_{ave}$, was estimated using the average flow rate and the average channel cross-sectional area, $A_{ave}$, found from the deformed and non-deformed cases.

$$u_{ave} = \frac{Q_{ave}}{A_{ave}} \tag{S14}$$

The velocity was integrated over time to determine the fluid displacement, $D_i$, for pore *i*.

$$D_i = \int_0^{t_{half}} u(t)dt \tag{S15}$$

$$\simeq u_{ave}t_{half}$$

### S2. Solute Concentration

#### S2.1. Pore Solute Concentration

The nutrient concentration, *C*, in each pore at any given time was determined using the iterative method outlined by Wang et al. and summarized below[4]. Mixing in the pores occurs as each half cycle causes an influx of fluid that spreads to evenly fill the pore. The initial tracer concentration in each pore, $C_{pi}^{n}$, is zero, where *n* indicates the cycle number and $p_i$ represents the pore number labeled consecutively based on increasing distance from the source channel. Concentrations are examined relative to the concentration in the source channel. The source channel, $C_{source}$, is assumed to always remain constant as the fluid is replenished as it flows and its dimensions are significantly larger than any side channels.

The volume of fluid influx during each half cycle, $q_i$, equals the product of the number of channels connecting pore $i$-1 to pore $i$, $N$, the average cross-sectional area of the channels and the fluid displacement.

$$q_i = NA_{ave}D_i \quad \text{(S16)}$$

Equation S16 represents the amount of tracer transported with the fluid influx, subtracting the fluid efflux volume for a given cycle. For simplicity, we can set the value of N to one without altering the analysis. The concentration of the first pore is

$$C_{p1}^{n} = C_{p1}^{n-1} + \frac{V_C C_{C1}^{n-1} + (q_1 - V_C)C_{SOURCE} - q_2 C_{p1}^{n-1}}{V + q_1 - q_2} \quad \text{(S17)}$$

where $V_c$ is the average channel volume over a half cycle and $V_p$ is the initial pore volume.

The concentrations in subsequent pores is given by:

$$C_{pi}^{n} = C_{pi}^{n-1} + \frac{V_C C_{Ci}^{n-1} + (q_i - V_C)C_{p(i-1)}^{n-1} - q_{i+1} C_{pi}^{n-1}}{V + q_i - q_{i+1}}. \quad \text{(S18)}$$

In most cases $q_i$ can be approximated as equal to $q_{i+1}$ so that the terms in the right-hand side of Equation S18 can be rewritten as follows:

$$C_{pi}^{n} = C_{pi}^{n-1} + \frac{V_C}{V} C_{Ci}^{n-1} + \frac{(q_i - V_C)}{V} C_{p(i-1)}^{n-1} - \frac{q_i}{V} C_{pi}^{n-1} \quad \text{(S19)}$$

At the end of each iteration, the value for the fluid concentration in the upstream and downstream channels, $C_{Ci}^{n}$ and $C_{C(i+1)}^{n}$, are updated accordingly.

$$C_{C1}^{n} = C_{SOURCE} \quad \text{(S20)}$$

$$C_{Ci}^{n} = C_{p(i-1)}^{n-1} \quad \text{(S21)}$$

$$C_{C(i+1)}^{n} = C_{pi}^{n-1} \quad \text{(S22)}$$

The Channel geometry, applied load magnitude and frequency were inputted into a MATLAB script to determine the channel-pore resistance of each segment and the resulting fluid displacement over a half cycle. The concentration equations S19-S22 were implemented iteratively with values for each pore outputted in an array.

If we combine terms in Equation S19 to the change in volume of the pore ($\Delta V$) and use the assumption that the concentration of solute in a channel is equal to that of the downstream pore we can derive the following equation for the change in solute concentration of a pore caused by one cycle of loading:

$$C_{pi}^{n}\text{-}C_{pi}^{n-1} = \frac{V_C}{V}\left(C_{Ci}^{n-1} - C_{p(i-1)}^{n-1}\right) + \frac{\Delta V}{V}\left(C_{p(i-1)}^{n-1} - C_{pi}^{n-1}\right). \quad \text{(S23)}$$

Equation S23 therefore separates out the effect of channel-pore geometry ($\frac{V_C}{V}$) from that of volumetric strain caused by mechanical loading ($\frac{\Delta V}{V}$).

### S2.2. Minimum Pore Nutrient Concentration Threshold

Bacteria sequestered inside a material require sufficient nutrients to remain viable. Bacteria require a carbon source, typically glucose, to sustain normal growth. Common growth media, such as lysogeny broth, is often supplemented with 0.1- 0.4% glucose to meet this need. The amount of glucose required by bacteria depends on several factors including the bacteria species and strain, temperature, and metabolic state of the bacteria. This analysis assumes that the initial population has no initial nutrient store. We estimate the nutrient needs of *E. coli* under standard growth conditions (37˚ C. and 200 rpm) using the findings from Seong et al. regarding nutrient metabolism in *E. coli* who found that, on average, *E. coli* requires 2.5 x $10^{-13}$ g/s of glucose to maintain viability. This corresponds to 65% of the glucose concentration in standard growth media (0.1% glucose)[5].

The glucose consumption rate for E. coli equals 0.4714 g glucose / g dry cell weight hr (gDCW h) [5]. The glucose required for the bacterial population in one pore is

$$\left(1.91\ x10^{-5}\,\frac{\text{gDCW}}{\text{pore}}\right)\left(0.4714\,\frac{\text{g glucose}}{\text{gDCW h}}\right)\left(\frac{1}{3600}\,\frac{\text{hr}}{\text{s}}\right) = 2.5\text{ x }10^{-9}\,\frac{\text{g glucose}}{\text{s pore}} \quad \text{(S25)}$$

The mass of bacteria in each pore can be found using a bacterial concentration of $10^{9}$ CFU/mL, an average pore volume for the mechanically loaded designs of 2.22 x $10^{-2}$ mL and an estimated cell mass of 8.6 x $10^{-4}$ gDCW [6].

$$\left(\frac{2.22x10^{-2}\text{ mL}}{\text{pore}}\right)\left(\frac{8.6x10^{-4}\text{ gDCW}}{\text{mL}}\right) = 1.91\ x10^{-5}\,\frac{\text{gDCW}}{\text{pore}} \quad \text{(S26)}$$

Assuming a loading frequency of 1 Hz, a loading magnitude of 60 kPa and an influx volume of 3.87 x $10^{-10}$ mL, the required concentration of the influx fluid is

$$\left(2.5 \text{ x } 10^{-9}\,\frac{\text{g glucose}}{\text{s}}\right)\left(\frac{1}{\frac{3.87\text{x}10^{-6}\text{mL}}{\text{s}}}\right) = 6.46\, x\, 10^{-4}\,\frac{\text{g glucose}}{\text{mL}} \qquad \text{(S27)}$$

The resulting concentration determined in (S24) can be compared to a source concentration of 0.1% glucose to find the minimum concentration relative to the source concentration.

$$\frac{6.46x10^{-4}\frac{\text{g glucose}}{\text{mL}}}{0.001\,\frac{\text{g}}{\text{mL}}}\; x\; 100\% = 64.5\% \qquad \text{(S28)}$$

This provides an estimate of the minimum pore concentration, relative to the source, required to keep resident bacteria in the same metabolic state observed by Seong et al. [5]. The minimum percentage concentration decreases to 16.1% if a 0.4% glucose source solution is used.

**S3. Diffusion**

The diffusion flux of the fluorescent dye from the main channel through the side channel and into the pores was estimated using Fick's first law of diffusion

$$J = -D\,\frac{dC}{dx} \qquad \text{(S29)}$$

where *D* is the diffusivity, *C* is the concentration and *x* is the position. The diffusivity of calcein was estimated to be 5 x $10^{-6}$ $cm^2/s$ [7]. This is similar to the value of fluorescein and other fluorescent tracers [8]. A source concentration of 1 mM Calcein was used [9]. It is important to note that particle size and solvent viscosity would affect diffusion, however these were not of concern as low concentrations were used to remain in the linear range of fluorescence.

An updated concentration could then be found based on the time, initial concentration, and geometry of the device. The transport due to diffusion was added to the iterative calculation when analyzing cyclic loading.

**S4. Bending**

Mechanical analysis of the PDMS device during cyclic loading was performed to ensure that bending of the device during loading did not greatly influence the optical plane. Any offset between the line of action of the force and the center of the device would generate a bending moment. The following estimate shows that deformation of the channels due to bending stress is negligible. The bending moment is

$$M = EIy \quad \text{(S30)}$$

where $I$ is the first moment area of Inertia for a rectangle and $y$ is the offset distance. The bending stress due to the maximum applied load, $P$, is given by:

$$\sigma_{bending} = \frac{My}{EI} \quad \text{(S31)}$$

$$= -\frac{Py^2}{EI}$$

$$= -\frac{(20\ \mathrm{N})(0.001\ \mathrm{m})^2}{(1.32x10^6\ \mathrm{Pa})\frac{(0.044\ m)(0.007\ \ \mathrm{m})^3}{12}}$$

$$= 1.20x10^{-2}\mathrm{Pa}$$

This value was negligible when compared with the estimated normal stress

$$\sigma_{normal} = PA \quad \text{(S32)}$$

$$= \frac{20N}{(0.044)(0.007)}$$

$$= 6.49x10^4\mathrm{Pa}$$

Therefore, bending could be neglected.
